# A Computational Method to Dissect Colonization Resistance of the Gut Microbiota against Pathogens

**DOI:** 10.1101/2022.01.06.475215

**Authors:** Shanlin Ke, Yandong Xiao, Scott T. Weiss, Xinhua Chen, Ciaran P. Kelly, Yang-Yu Liu

## Abstract

The indigenous gut microbes have co-evolved with their hosts for millions of years. Those gut microbes play a crucial role in host health and disease. In particular, they protect the host against incursion by exogenous and often harmful microorganisms, a mechanism known as colonization resistance (CR). Yet, identifying the exact microbes responsible for the gut microbiota-mediated CR against a particular pathogen remains a fundamental challenge in microbiome research. Here, we develop a computational method --- Generalized Microbe-Phenotype Triangulation (GMPT) to systematically identify causal microbes that directly influence the microbiota-mediated CR against a pathogen. We systematically validate GMPT using a classical population dynamics model in community ecology, and then apply it to microbiome data from two mouse studies on *C. difficile* infection. The developed method will not only significantly advance our understanding of CR mechanisms but also pave the way for the rational design of microbiome-based therapies for preventing and treating enteric infections.

## Introduction

Rather than simple passengers in and on our bodies, commensal microorganisms play key roles in human physiology and diseases^1^. In particular, the gut microbiota of healthy individuals provides colonization resistance (CR) against pathogens through multiple mechanisms, including secretion of antimicrobial products, nutrient competition, support of epithelial barrier integrity, bacteriophage deployment, and immune activation^2^. Yet, identifying the exact microbial species responsible for the gut microbiota-mediated CR against a particular pathogen, i.e., identifying causal species that directly inhibit the growth of the pathogen, remains a fundamental challenge in microbiome research^3–9^.

Microbial interactions can be mediated by direct secretion of substances such as bacteriocins^10,11^, ecological competition between the microbes^12^, metabolite exchange^13^, or/and immune system modulation^14–16^. Note that the microbial interactions discussed here are fundamentally different from associations or co-occurrences calculated from similarity-based techniques^17^. The former can be used to predict the dynamic behavior of microbial communities, while the latter cannot. Various methods have been developed to infer microbial interactions and map the ecological network of microbial communities. Yet, those methods require either high-quality time-series data^18,19^ or a large number of steady-state samples^20^, which significantly limit their practical application.

For host-associated microbiome, the standard approach for identifying phenotype-associated microbes has been the microbe-wide association study (MWAS), which is analogous to genome-wide association studies that are commonly used to identify genetic variants linked to a given phenotype. In a standard MWAS, microbial compositions are compared between two populations of interest (e.g., diseased and healthy), and differentially abundant microbes are identified to correlate with the phenotype of interest^21^. However, due to the extensive complexity of the human microbiota, particularly the microbial interactions, MWAS typically generates a long list of commensals implicated as biomarkers of disease, with no clear relevance to disease pathogenesis. Identifying causal microbes using MWAS has remained difficult.

Recently, the so-called microbe-phenotype triangulation (MPT) method was developed to identify causal microbes that are likely to influence disease pathogenesis^22^. This method compares the microbial communities of groups of hosts that elicit divergent phenotypes (e.g., susceptibility to disease) to pinpoint causal microbes. While comparisons between two phenotype groups (i.e., MWAS) often yields a long list of differentially abundant microbes, comparing multiple phenotype groups and checking their overlap might reduce this list down to a small number of microbes. For example, MPT has been applied to identify microbes that may protect mice from a chemically induced colitis^22^. The key assumption of the MPT method is that if any taxa were truly relevant to disease pathogenesis, they would be present in all pair-wise comparisons of different phenotype groups. Yet, this key assumption has not been theoretically justified. Species/strains from these causal taxa (if they exist, typically at a very high taxonomic level, e.g., family) have to be tested individually to determine whether they play any causal role in determining the observed host phenotype. In fact, the whole MPT framework is quite empirical, and lacks rigorous justification from the community ecology perspective.

In this work, we propose a generalized microbe-phenotype triangulation (GMPT) method to systematically identify microbes that mediate CR against pathogens. GMPT is based on the following key hypothesis: those differentially abundant species that appear in most of the pair-wise phenotype comparisons and whose abundances display strong negative (or positive) correlation with the abundance of a pathogen are preventive (or permissive) causal species that directly inhibit (or promote) the growth of the pathogen. Unlike the original MPT method, GMPT can be theoretically validated. In this work, we systematically validated GMPT using a classical population dynamics model in community ecology. Then we applied it to microbiome data from two mouse studies on *C. difficile* infection (CDI), identifying a comprehensive list of preventive and permissive microbial species that may be directly related to the pathogenesis of *C. difficile*. The presented method sheds light on the rational design of microbiome-based therapies for preventing and treating CDI and other infections.

## Results

### Workflow of GMPT

An overview of the GMPT method is shown in Fig.1. *First*, we divide microbiome samples into different phenotype groups based on the pathogen abundance or disease severity (Fig.1a,b). *Second*, we perform differential abundance analysis for each phenotype pair to get a pool of differentially abundant species using standard tools, e.g., ALDEx2^23^. We then rank those differentially abundant species based on their frequency present in all pair-wise phenotype comparisons in a descending order (Fig.1c). If there is a tie, we will further rank species based on their differentiability in a descending order. We use effect size (relative fold difference) to quantify species’ differentiability in ALDEx2. *Third*, for each differentially abundant species, we calculate the Spearman correlation (denoted as *ρ*) between its mean relative abundance and the ascending order of the different phenotype groups (ranked based on their ability to confer CR against pathogen, which can be quantified by the pathogen abundance). Finally, those differentially abundant species with positive (or negative) *ρ* are candidate causal species that can directly promote (or inhibit) the growth of pathogen, respectively. Notably, our GMPT pipeline did not use any threshold to suggest causal species. Instead, we offer a ranking of their candidacy (based on their occurrence in all pair-wise DAAs).

**Fig. 1.**
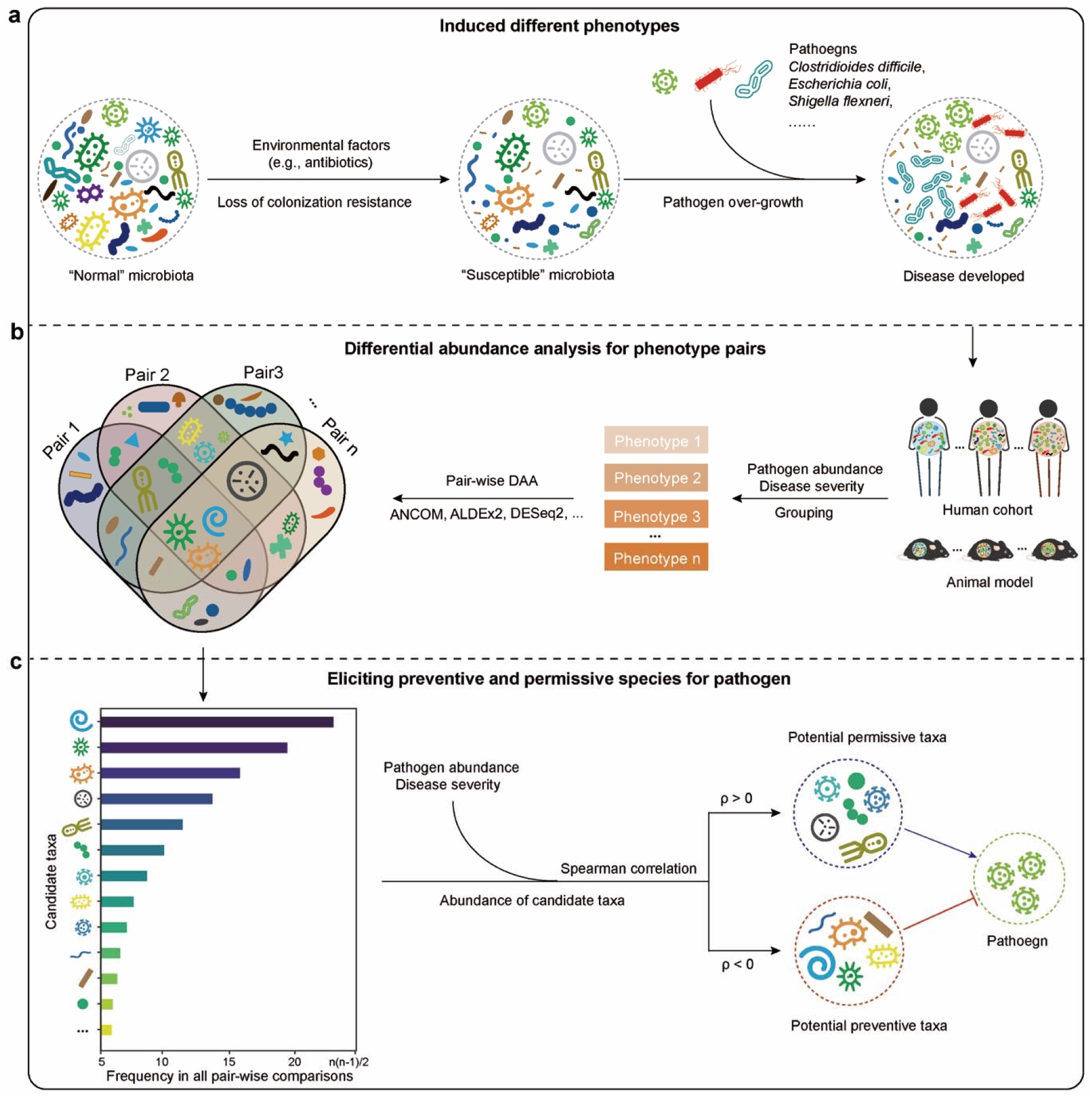
Summary of overall workflow. **a**. The gut microbiome is critical in providing resistance against colonization by exogenous microorganisms. Changes in microbiota composition, and potential subsequent disruption of CR, can be caused by various environmental factors such as diet or antibiotics, thereby providing opportunities for pathogens to colonize the intestines and ultimately cause disease. **b**. The microbial samples from a human cohort or animal model can be grouped based on the phenotype (resistance ability against colonization of pathogen). Differential abundance analysis will be carried out on each possible pairwise comparison. The differentially abundant microbes pool comes from each pairwise analysis and the Venn diagram shows the frequency distribution of differentially abundant microbes among all the comparisons. **c.** The differentially abundant taxa will be ranked based on their frequency appearing in all the pairwise comparisons and differentiability (descending order). We further use the average relative abundance of species and ascending rank index of the ability to confer colonization resistance against pathogen (e.g., the abundance of specific pathogen or the outcome of host) in all phenotypes to calculate the Spearman correlation coefficient. Here positive (negative) Spearman correlation coefficient (ρ) represent the permissive (preventive) taxa of pathogen. And the spearman correlation with 0 means the taxa may be neutral to the growth of pathogen.

An effective way to investigate the effect of the gut microbiota on a specific phenotype is co-housing of affected and unaffected animals^24^. Due to coprophagy, during cohousing animals may feed on feces or ingest feces by self-grooming. A previous study demonstrated that co-housing mice susceptible to chemically induced colitis with resistant mice protected the susceptible mice, but also converted resistant mice to be more susceptible^25^. In other words, co-housing naturally leads to intermediate phenotypes. This phenomenon inspires us to systematically validate our GMPT method *in silico*.

### Validate GMPT using a community ecology model

We model the gut microbiota of different hosts as different local communities assembled from a global species pool (or metacommunity) of *N* species governed by a universal set of dynamical rules^26^. In particular, we assume the population dynamics of any local community can be described by a set of ordinary differential equations (ODEs):

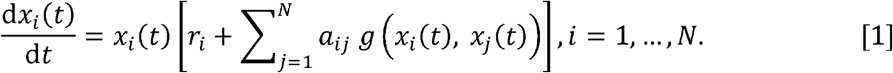

Here, *x_i_*(*t*) represents the abundance of species-*i* at time *t*, 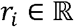 is the intrinsic growth rate of species-*i*, 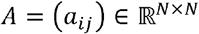 is the species interaction matrix, and 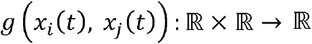 is the so-called *functional response* in community ecology, which models the intake rate of a consumer as a function of food density. A linear functional response *g*(*x_i_, x_j_*) = *x_j_* corresponds to the classical Generalized Lotka-Volterra (GLV) model, which has been used in many existing ecological modeling works of host-associated microbial communities ^18,20,27–32^. In this work, we will also use the GLV model. The species interaction matrix *A* encodes the ecological network 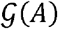: there is a directed edge 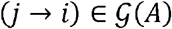 if and only if *a_i□_* ≠ 0. In GLV model, *a_ij_* >0 (< 0) means that species-*j* promotes (inhibits) the growth of species-*i*. The ecological network of a small community is shown in Fig.2a, where the edge thickness corresponds to the absolute value of *a_ij_*, i.e., the interaction strength. In this work, we use the GLV model to generate synthetic data to validate the GMPT method. In particular, we want to identify those species that directly promote or inhibit the growth of a pathogen.

**Fig. 2.**
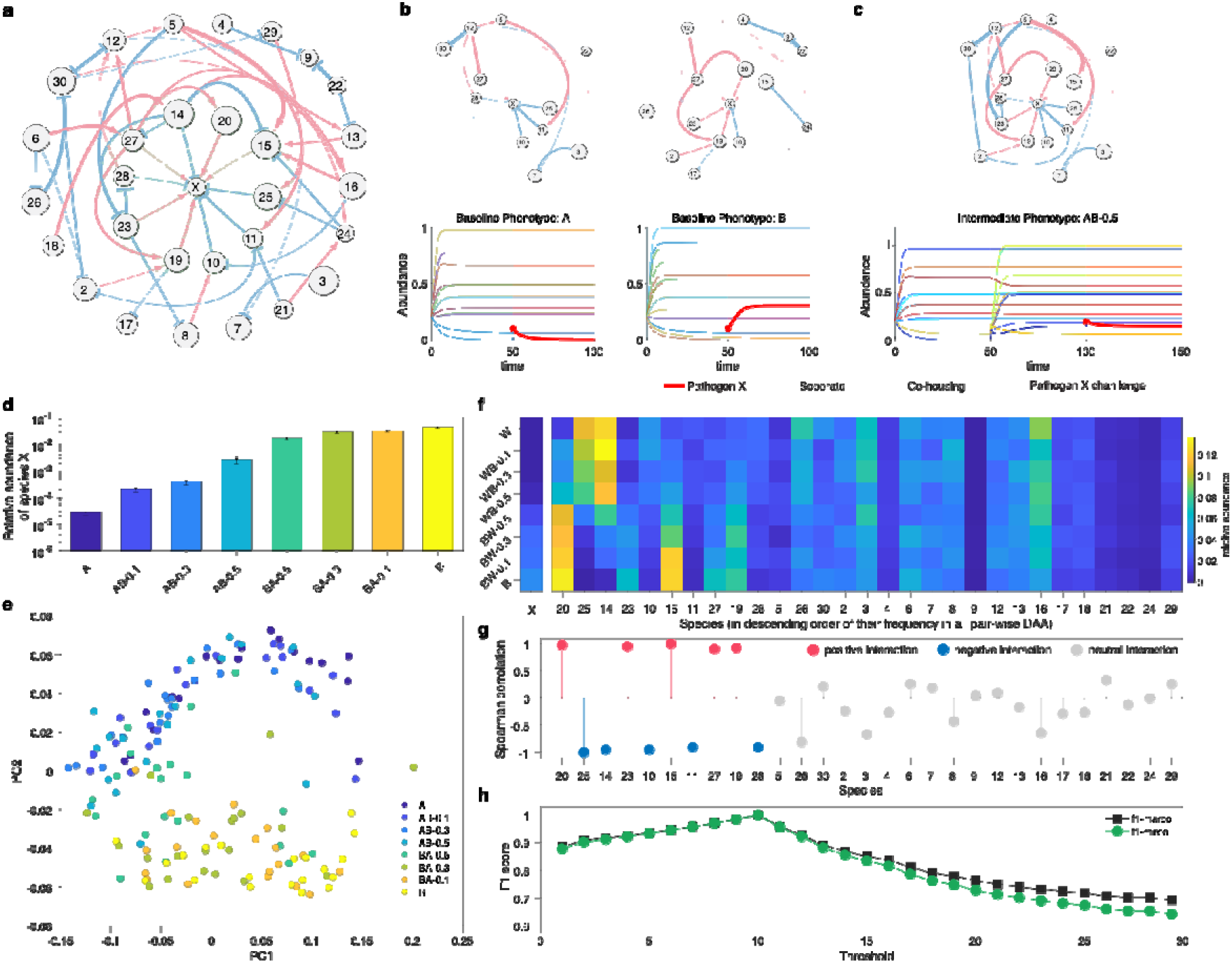
The ecological modeling framework of GMPT. **a.** A synthetic ecological network with 30 species and node X represents the specific pathogen. The blue (or red) edge means the pairwise inhibition (or promotion) effect of one species on another species. The thickness of edges corresponds to the absolute values of interaction strength. The green area covers the species with directed inhibition or promotion interactions on pathogen X. In this small synthetic community, GMPT aims to identify these species with one-step causal effect on pathogen X. **b.** The temporal abundance and corresponding ecological networks of two samples from two baseline phenotypes A and B, respectively. The grey nodes indicate the absent species in phenotype A or B. After pathogen X challenge, it will not grow up in A subject but bloom in B subject. **c**. Our modeling framework mimics the co-housing process to generate the intermediate phenotype AB-0.5 that a sample from phenotype A was co-housed with another sample from phenotype B, and there was species from B are transferred to A. After co-housing, the original subject will transfer from phenotype A to AB-0.5 because pathogen X will not be suppressed to extinction in the new phenotype. **d.** Relative abundance of pathogen of 2 baseline phenotypes and 6 co-housing phenotypes. The error bar represents the standard error of the mean (SEM) calculated from 20 independent simulations for each phenotype. **e.** PCoA of 8 phenotypes. The Bray-Curtis dissimilarity is used in the PCoA throughout this manuscript. **f.** Abundance distribution of microbial species across different phenotypes. For each phenotype pair, we apply ALDEx2 to identify those differentially abundant species. The-axis corresponds to the species in a descending order of their present frequency in all pair-wise DAA. **g.** Then we calculate the spearman correlation of species (except pathogen X) between two vectors ***p*** = (*p*_1_,…, *p_m_*) and ***q*** = (*q*_1_,…, *q_m_*) where *p_a_* is the average relative abundance of the pathogen in phenotype-*a*, and *q_a_* is the average relative abundance of species *i* in phenotype-*a*. Blue (red or grey) indicates the inhibition (promotion or neutral, respectively) effects of other species on pathogen X. **h.** F1-score of three-way classification (promotion, inhibition and neutral) under different top-*K*. The *K* value in *x*-axis is sorted by the descending present frequency.

We first generate the two baseline phenotypes (denoted as “A” and “B”) from a microbial community with 30 species (see Fig. 2a for the ecological network, where the center node X represents the pathogen that we want to decolonize, and those species directly interacting with X are highlighted in a green circle). The two baseline phenotypes are selected based on the criterion that the pathogen cannot grow up in A but will bloom in B (see Fig.2b for the simulated temporal behavior of species abundances before and after the pathogen challenge, where the abundance of the pathogen is highlighted in red). The detailed procedure of generating baseline phenotypes is shown in Supplementary information (SI Sec.1&2).

Then we generate intermediate phenotypes by simulating the “co-housing” process. In particular, we transfer a certain fraction of species from one community to another, assuming the fraction of transferred species is proportional to the co-housing time. For example, AB-*x* means that a community/sample from phenotype A was co-housed with a community/sample from phenotype B, and a fraction *x* of species from B are transferred to A. Finally, the community evolves to a new intermediate phenotype, denoted as AB-*x*. Note that, AB-*x* is different with BA-*x*. Fig. 2c displayed the temporal behavior of a community from phenotype A in separate housing, co-housing, and pathogen challenge. Note that since *x* = 0.5, 50% of species from B are transferred to A, rendering a more susceptible phenotype, where the pathogen can successfully colonize with considerable abundance (higher than that of A, but lower than that of B, hence representing an intermediate phenotype). By tuning *x*, we can generate various intermediate phenotypes. Fig.2d showed the different phenotypes ranked by their average relative abundance of the pathogen in an ascending order. Fig.2e illustrated the compositional difference of communities in different phenotypes in the PCoA plot.

Suppose we generate in total *m* phenotypes (including the two baseline phenotypes). Then we perform differential abundance analysis for all the *m*(*m* – 1)/2 phenotype pairs, e.g., using ALDEx2, to identify differentially abundant species. We rank those differentially abundant species (except the pathogen X) based on their number of occurrences in the pair-wise differential abundance analyses (DAA). Then for each of the differentially abundant species *i* (except pathogen X), we calculate the Spearman correlation *ρ_i_* between two vectors ***P*** = (*p_i_*,…, *p_m_*) and ***q*** = (*q*_1_,…, *q_m_*), where *p_a_* is the average relative abundance of the pathogen in phenotype-*a*, and *q_a_* is the average relative abundance of species *i* in phenotype-*a*. The positive (or negative) Spearman correlation *ρ_i_* indicates that species-*i* tends to have higher (or lower) abundance in a phenotype with higher abundance of the pathogen. Besides, for those species that directly interact with pathogen X should be identified as differential abundant species by DAA, and they should appear more in all pair-wise DAAs than those species that do not directly interact with pathogen X.

Our key hypothesis is that those differentially abundant species with positive (or negative) *ρ_i_* and higher occurrence in the phenotype pair comparisons tend to have a higher probability to promote (or inhibit) the growth of the pathogen. Interestingly, this hypothesis was validated in the small synthetic microbial community shown in Fig.2a. For example, the vectors ***p*** and ***q*** for pathogen X and other species are shown as columns in the heatmap of Fig. 2f, respectively. The rows in Fig.2f were shown in a descending order of the species’ occurrence in all pair-wise DAAs. The Spearman correlation *ρ_i_* can be easily calculated from the heatmap. The results are shown in Fig.2g. We found that: (i) those species that directly interact with the pathogen X appear more in pair-wise DAAs than those species that do not directly interact with X; (ii) those species that directly promote (or inhibit) the growth of pathogen X have positive (or negative) *ρ_i_*. The results demonstrated that combining the occurrence in all pair-wise DAAs and Spearman correlation can indeed identify those species that directly promote or inhibit the growth of the pathogen.

Note that for synthetic data we know the ground truth. Hence, we can quantify the performance of GMPT based on the F1-score of the three-way classification using the top-*K* candidate causal species. (See SI Sec.3 for details on the three-way classification.) We found that the F1-score reaches its maximum when *K* equals the number of species that directly interact with the pathogen. Hence the maximum of the F1-score can be used as an evaluation metric of the GMPT performance.

To systematically evaluate the performance of GMPT, we generated synthetic data using the GLV model with different levels of network connectivity (*C*), characteristics inter-species interaction strength (*δ*), and measurement noise of species abundance (*η*), as well as different number of phenotypes and different sample size. We found that in general GMPT’s performance stabilizes when the number of phenotypes is larger than 4, and the performance increases with increasing sample size (Fig.3). Interestingly, GMPT’s performance is not very sensitive to other model parameters *C*, *δ*, and *η*.

**Fig. 3.**
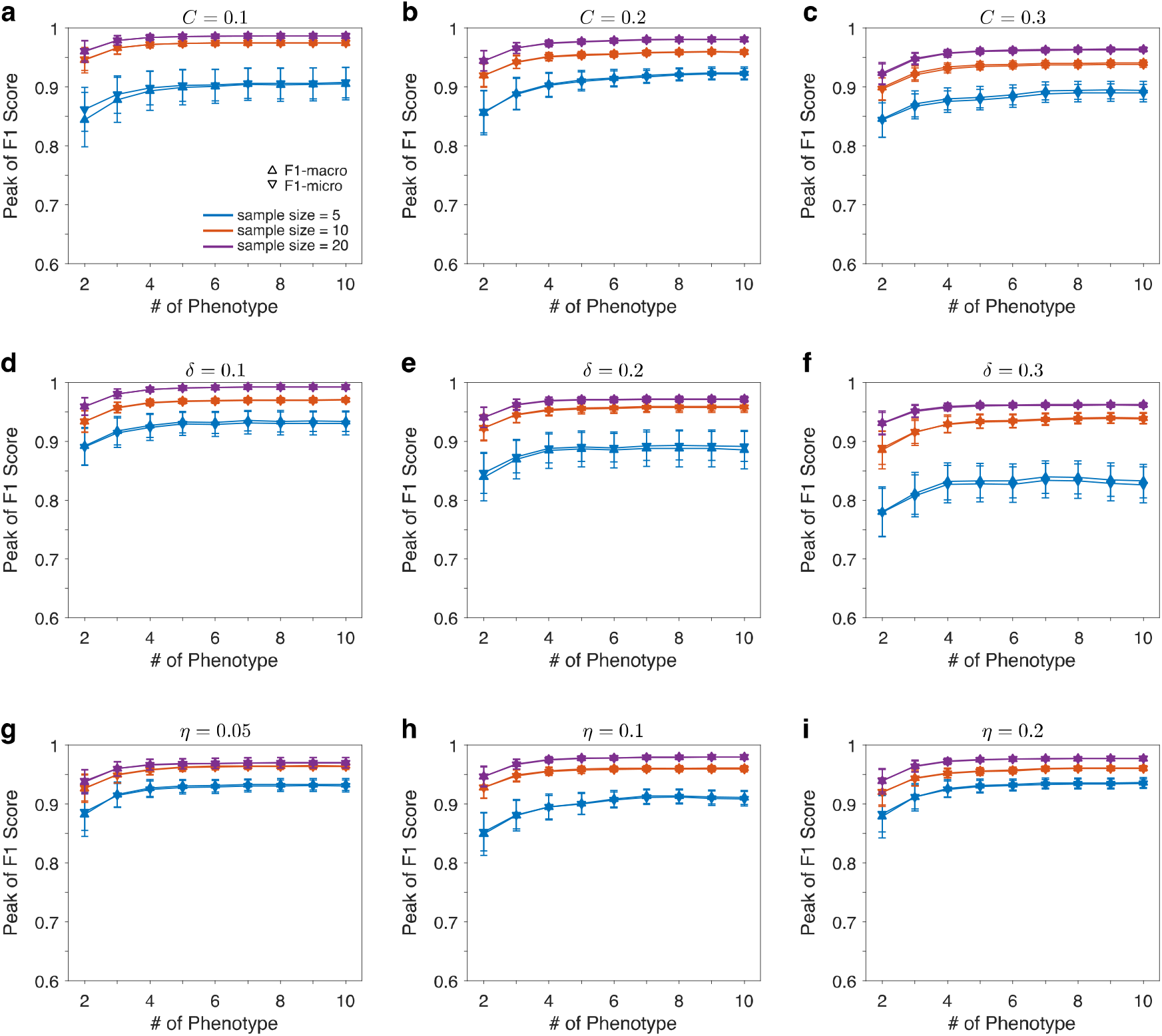
The systematical evaluation of GMPT to identify the causal species on a particular pathogen under different parameters. The peak of F1-score (macro- and micro-average) with different number of sample size and phenotypes when the network connectivity (denotes *C* in panels **a-c**), standard variance of *a_ij_* (*δ* in panels **d-f**) and measurement error of species abundance (*η* in panels **g-i**) take different values. The *x*-axis means the number of 10 phenotypes used in GMPT: A, B, AB-0.1, AB-0.2, AB-0.3, AB-0.5, AW-0.1, AW-0.2, AW-0.3, AW-0.5. The network connectivity, *C*, is the probability that there will be a directed edge between any two taxa. The standard variance of *a_ij_*, *δ*, takes from 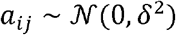. The measurement noise of species abundance, *η*, takes *x_i_* = *x_i_* + *ηU*[–*x_i_*, *x_j_*]. In panels a-c, the simulations fixed *δ* = 0.2, *η* = 0.05, and changed *C* = 0.1,0.2,0.3. In panels d-e, the simulations fixed *C* = 0.2, *η* = 0.05, and varied *δ* = 0.1,0.2,0.3. In panels g-i, the simulations fixed *C* = 0.2, *δ* = 0.2, and varied *η* = 0.05,0.1,0.2. The error bar represents the standard error of the mean (SEM) calculated from 10 independent ecological networks.

Numerous methods have been developed to identify differentially abundant species from the comparison of case and control microbiome samples^33^. To further validate the robustness of GMPT across different DAA methods, we applied ANCOM^34^ in GMPT pipeline. In ANCOM, a species’ differentiability can be quantified by its mean W-score, and a high mean W-score indicates the greater likelihood that the null hypothesis can be rejected, indicating the number of times a parameter is significantly different between groups. Interestingly, similar results were observed in GMPT using ANCOM (Figs. S1-S2).

### Real data application

To illustrate the proposed methodology, we firstly applied it to the murine gut microbial samples from a study of antibiotic-induced effects on CR against *C. difficile*^35^. Numerous studies have confirmed that the ability of microbiota to protect against pathogens (e.g., *C. difficile*) can be severely impaired by antibiotic treatment^16,35^. This dataset was a combination of multiple different perturbations (e.g., different antibiotic classes, doses, and recovery time), that allowed us to generate distinct community profiles that displayed a range of susceptibilities to *C. difficile* colonization. We then applied an additional filtering step to only obtain the samples with *C. difficile* challenges. A total of 132 samples from 132 mice were analyzed. We first split this cohort into 8 groups based on the levels of *C. difficile* colonization (Methods) in mice (Fig. 4a,b). The PCoA plot also indicated a distinct microbial structure (PERMANOVA test, *P*-value = 0.0001, Bray-Curtis dissimilarity) among different groups (Fig. 4c). We then identified differentially abundant bacteria in all twenty-eight (all possible combination from 8 groups) pairwise comparisons using ALDEx2. Given the difference underlying microbial sources for these mice, all pairwise analysis yielded a taxa pool of 119 differentially abundant ASVs presented at least one pairwise comparison. And the maximum frequency of differentially abundant ASVs presented in the pairwise comparisons was 11, and only three ASVs (ASV5: *Muribaculum intestinale*; ASV22: *Eubacterium ventriosum*; ASV60: Unclassified Clostridiales) were identified from those 11 pairwise comparisons. It has been suggested that CR against *C. difficile* was never attributable to a single microbe, but rather it was the results of multiple interacted microbial species. The top 30 candidate promoters or inhibitors were showed in Fig. 4d and Table S1.

**Fig. 4.**
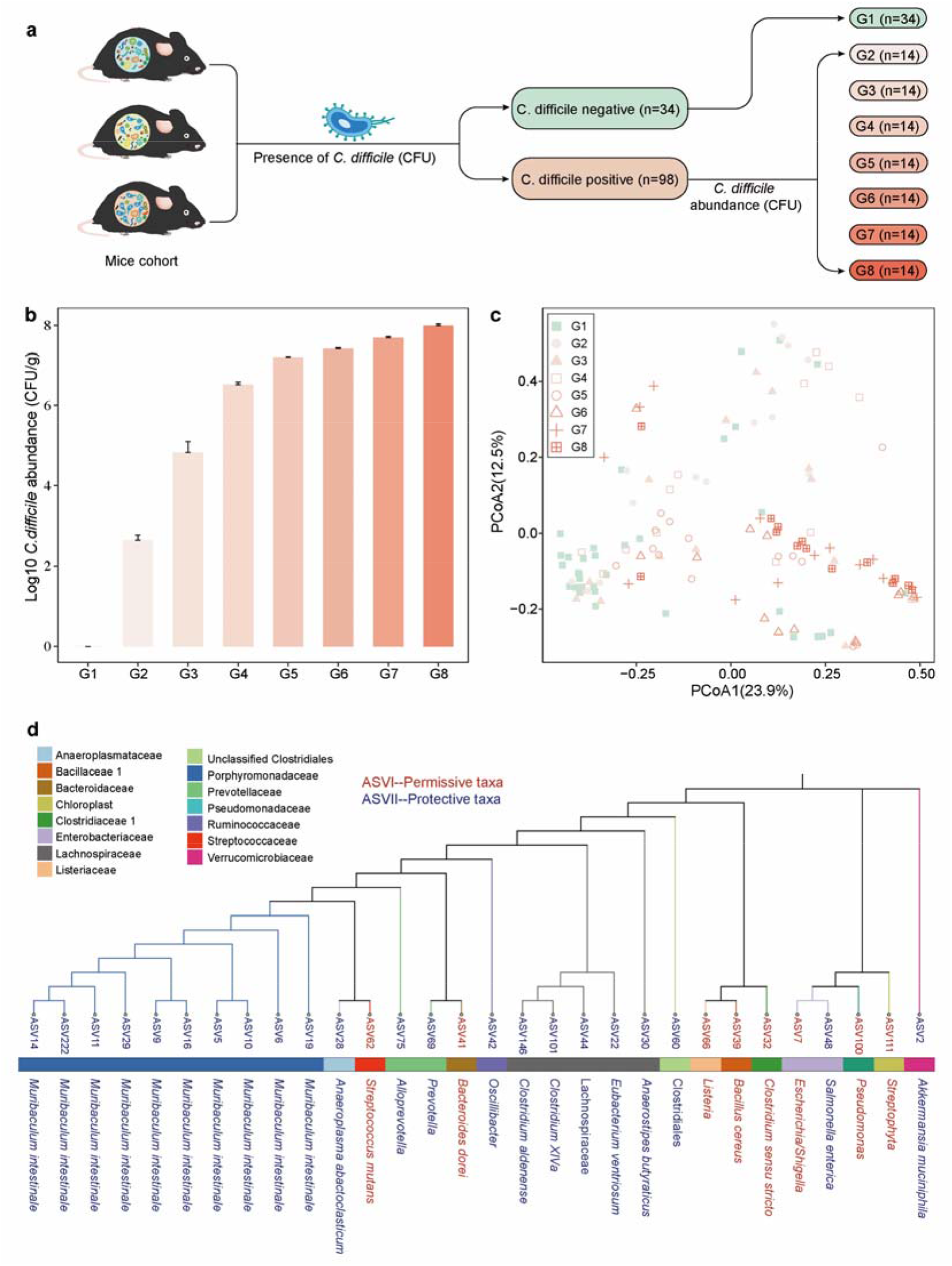
The *C. difficile* infection related taxa identified by GMPT in dataset I. **a.** The flowchart for obtaining different phenotype groups. **b.** Comparison of the abundance of *C. difficile* (CFU/g feces) in each group. Bars represents the values of mean ± s.e.m. **c.** PCoA plot of fecal microbiota in mice, indicating heterogeneity in gut microbial communities induced by antibiotics. **d.** A phylogenetic tree showing the *C.difficile* infection realted ASVs identified using GMPT. The colors of horizontal bar represents family level and the different colors of ASV and taxonomy information represents permissive or protective taxa.

Prior studies between the gut microbiome and diseases often yielded mixed results, which is a classical problem in the microbiome field. To further demonstrate the advantage of our method, we analyzed the second dataset related to gut microbiota and *C. difficile* colonization^36^ and then compared the findings with the first dataset. The original study aimed to elucidate how the gut bacterial community changes concurrently with clearance of *C. difficile* colonization. Based on the status of *C. difficile* colonization (i.e., not colonized, cleared, and colonized), we grouped this cohort into 7 groups (SI, Fig. 5a, b). As illustrated in the Fig. 5c, PCoA results indicated that the fecal samples from different phenotypes were clustered differently (PERMANOVA test, *P*-value = 0.0001, Bray-Curtis dissimilarity). Following our pipeline, all pairwise analysis (21 pairwise comparison from 7 groups) yielded a taxa pool of 153 differentially abundant ASVs presented at least one pairwise comparison. The top 30 candidate promoters or inhibitors of *C. difficile* are shown in Fig. 5d and Table S2.

**Fig. 5.**
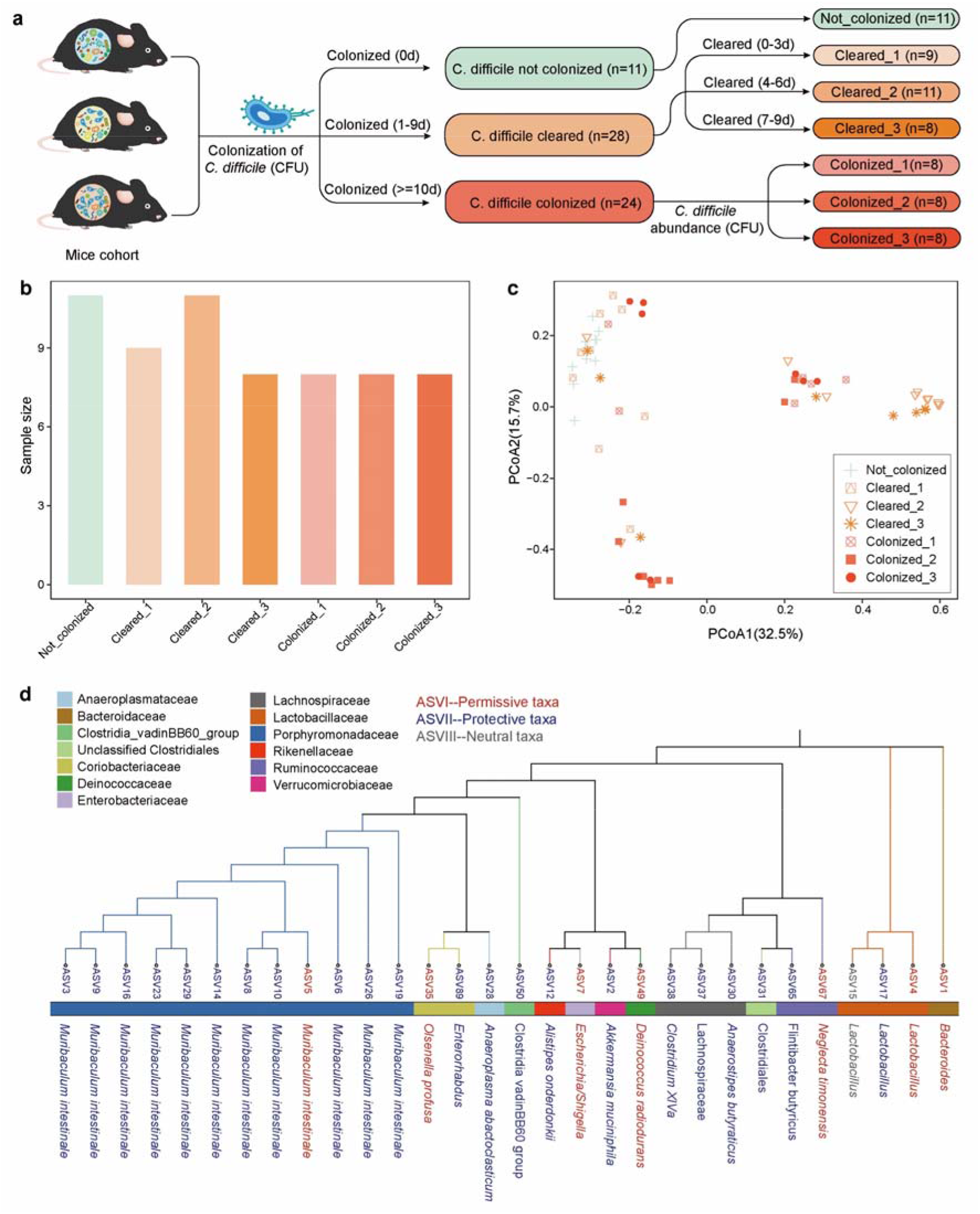
The *C. difficile* infection related taxa identified by GMPT in dataset II. **a.** The flowchart for obtaining different phenotype groups. **b.** Comparison of the abundance of *C. difficile* (CFU/g feces) in each group. Bars represents the values of mean ± s.e.m. **c.** PCoA plot of fecal microbiota in mice, indicating heterogeneity in gut microbial communities induced by antibiotics. **d.** A phylogenetic tree showing the *C.difficile* infection realted ASVs identified using GMPT. The colors of the horizontal bar represents family level and the different colors of ASV and taxonomy information represents permissive, protective or netural taxa.

Importantly, we found 40% (12/30, top 30) overlapped ASVs were identified from two datasets. Among these 12 overlapped taxa, most of (91.67%, 11/12) ASVs showed a consistent relationship with *C. difficile* (Fig. S3, Table S1, and Table S2). This provides evidence that the GMPT pipeline can robustly identify the disease-related microbial features across studies with similar designs. The conflicting ASVs between the two datasets could be explained by the labeling, number of phenotypes and sample size in each phenotype. Among these 11 candidate casual bacteria, we identified 10 preventative ASVs (i.e., *Muribaculum intestinale*, Anaerostipes butyraticus, *Akkermansia muciniphila*, *and Anaeroplasma abactoclasticum*) and 1 permissive ASVs (i.e., *Escherichia/Shigella*). Previous studies also indicated that *Muribaculum intestinale^31^*, Anaerostipes^38–40^, and *Akkermansia muciniphila*^4,4^ decreased in *C. difficile-positive* subjects or inhibited the growth of *C. difficile*. It has been shown that *Escherichia/Shigella* was significantly enriched in *C. difficile-positive* subjects or co-infected with *C. difficile*^43–45^. These results demonstrate the ability of GMPT to streamline the discovery of microbes that are associated with colonization resistance.

To further validate the robustness of GMPT across different differential abundance analysis tools in real data, we also compared the results of GMPT using ALDEx2^23^ and ANCOM^34^. As shown in Fig. S4a, GMPT identifies highly overlapped candidate lists between ALDEx2 and ANCOM at both dataset I (80.00%, 24/30) and dataset II (90.00%, 27/30). Moreover, 53.33% (16/30, top 30) overlapped ASVs were identified from two datasets using ANCOM (Fig. S4b). Interestingly, the overlapped ASVs between dataset I and dataset II identified using ANCOM and ALDEx2 also showed highly consistent (Fig. S4). This result demonstrates the robustness of GMPT to identify microbes that potentially inhibit (or promote) the growth of the pathogen.

## Discussion

Although numerous studies have described the composition of the intestinal microbiota in subjects with specific pathogen colonization or infection, specific alterations associated with loss of colonization resistance and subsequent development of disease from an ecological point of view remain unclear. To move beyond the basic associations, we employed a classical GLV model to systematically justify the methodology of GMPT. In both simulation and experimental datasets, GMPT presents robust and improved performance for inferring the candidate inhibitors and promoters in infectious diseases caused by pathogens.

Through our ecological modeling framework, we were able to generate many different phenotypes (CR) related to specific pathogen (e.g., *C. difficile*) infection. Our approach is based on a clearly defined assumption that if any taxa were truly related to CR, they would be present in most of the pairwise differential abundance comparisons. We reasoned that the causal agent for CR should be a small group of bacteria rather than a specific microorganism. Using this model-based approach to estimate the pathogen related microbiota, we could estimate the relative contributions of taxonomically diverse bacteria to the CR against pathogen. Moreover, our modeling framework clearly demonstrated the effects of number of sample size and phenotypes on the performance of our pipeline. These findings may guide researchers in the experimental design of microbiome related studies.

Antibiotic treatments (e.g., antibiotic classes, doses, and recovery periods) are able to perturb the gut microbiota of the host to alternative dysbiotic states^46–48^, which allow us to obtain different phenotypes of pathogen colonization. In the CDI datasets, we identified a number of ASVs associated with the pathogenesis of CDI, and most of those top candidate promoters or inhibitors have also been reported as potential associations with CDI by multiple earlier studies. Interestingly, GMPT also reliably identified several protective species (e.g., taxa associated with the Porphyromonadaceae (*Muribaculum intestinale*) and *Alistipes*) and permissive species (e.g., *Escherichia*), which have been reported previously in the original studies^35,36^. Finally, GMPT was able to identify a comprehensive list of gut microbiota that can mediate CR against *C. difficile*, where some of those microbes were missed in the original study^35,36^. Moreover, the overlapping results between different differential abundance analysis tools may help to further narrow down the potential candidate species list. Although this study focuses on *C. difficile* infection as an example, general principles regarding CR mediated by the gut microbiome can also be applied to pathogens such as bacteria (e.g., *Escherichia coli* and *Salmonella*), viruses (e.g., HIV and SARS-CoV-2), and fungi (*Candida auris* and *Aspergillosis*).

In light of these findings, the current methodology still has limitations. First, based on our previous studies on the origins and control of community types in human microbiome^29^ and simulation process of FMT^28^, we treated the stochastic effects as negligible in our current modeling framework. In general, we can consider stochastic effects in our modeling framework but we believe this will not fundamentally affect our co-housing process. Second, the degree of net impact of a species on a specific pathogen is mainly based on the GLV model (only considers linear functional response and pair-wise microbial interactions). The way to systematically calculate the net impact in a complicated population model with non-linear functional response or higher-order interactions remain challenge. Third, we did not provide an exact threshold to select those permissive or preventive species from a clinical translational perspective. Because the community colonization resistance should be considered as byproduct of an assemblage of microbial populations rather than a specific population of microbe. We need to consider not only the dominant effect species, but also the small effect species as much as possible. However, we suggest that users can primarily focus on the overlapped results identified from different studies or differential abundance analysis tools. Furthermore, additional experiments are needed to assess the casual mechanisms of candidate species from an ecological perspective.

In conclusion, we theoretically justified the GMPT method by a classical population dynamics model in community ecology. Our results demonstrated that the GMPT pipeline is a powerful tool to identify the relative contributions of taxonomically diverse microbes to the CR against specific pathogen and their determinants in the disease pathogenesis. The method presented here paves the way to rationally designing of combination of microbes, which may be translated to better preventive and therapeutic approaches tailored to specific pathogen-induced diseases.

## Data and code availability

The sequencing data from the first CDI dataset^35^ are publicly available with SRA accession number SRP057386 (https://www.ncbi.nlm.nih.gov/sra/?term=SRP057386). The sequencing data from the second CDI dataset^36^ can be found at: https://www.ncbi.nlm.nih.gov/bioproject/PRJNA674858. Analysis code is available via: https://github.com/xiaoyandong08/GMPT.

## Author Contributions

Y-Y.L conceived and designed the project. Y.-Y.L and Y.X. developed the computational method. S.K. analyzed all the real data. Y.X. did all the numerical calculations. S.K., Y.X. and Y.-Y.L. wrote the manuscript. S.T.W, X.C, and C.P. K reviewed and edited the manuscript. All authors approved the manuscript.

## Acknowledgement

Y.-Y.L. acknowledges grants from National Institutes of Health (R01AI141529, R01HD093761, RF1AG067744, UH3OD023268, U19AI095219 and U01HL089856). Y.X. acknowledges funding support from National Natural Science Foundation of China (Grant No. 61902418).

